# Whole genome sequencing of 76 Mexican Indigenous reveals recent selection signatures linked to pathogens and diet adaptation

**DOI:** 10.1101/2024.07.26.605344

**Authors:** F. Miron-Toruno, E. Morett, I. Aguilar-Ordonez, A.W. Reynolds

## Abstract

Whole genome scans for natural selection signatures across Mexican Indigenous (MI) Populations remain underrepresented in the literature. Here, we conducted the first comparative analysis of genetic adaptation in MI populations using whole genome sequencing (WGS) data from 76 individuals representing 27 different ethnic groups in Mexico. We divided the cohort into Northern, Central, and Southern populations and identified signals of natural selection within and across populations. We find evidence of adaptation to pathogenic environments in all our populations, including significant signatures in the Duffy blood group gene in Central MI populations. Despite each region exhibiting unique local adaptation profiles, selection signatures on *ARHGAP15, VGLL4, LINGO2, SYNDIG1*, and *TFAP2B* were common to all populations. Our results also suggest that selection signatures falling within enhancers or promoters are usually connected to non-coding features, with notable exceptions like *ARHGAP15* and *GTDC1*. This paper provides new evidence on the selection landscape of Mexican Indigenous Populations and lays the foundation for additional work on Mexican phenotypic characterization.

**Significance statement:** Previous research has identified distinct patterns of genomic adaptation across the different regions of Mexico, highlighting evidence of natural selection within metabolic and immune-related genes. However, the characterization of the Mexican selection landscape from a whole-genome perspective remains unexplored. Here, we conducted the first whole-genome scan for natural selection in 76 Mexican Indigenous individuals from 27 different ethnic groups divided into Northern, Central, and Southern populations. Our findings revealed distinct local adaptation profiles for each Mexican region, with different evidence of adaptation to pathogenic environments across these groups. In contrast, all populations had common selection signatures on *ARHGAP15, VGLL4, LINGO2, SYNDIG1*, and *TFAP2B*. This paper provides new evidence on the genetic basis of adaptation of Indigenous groups in Mexico. Moreover, it provides a foundation for additional work on Mexican phenotypic characterization.

## Introduction

The genomes of Mexican Indigenous (MI) populations offer a window into the evolutionary history and adaptation of humans in the Americas. Recent archeological and genetic findings suggest that modern humans arrived on the American continent during the late Pleistocene period (1,2). This initial wave of migration, characterized by its wide and rapid expansion, brought human populations to new and often extreme environments that imposed strong selective pressures (2,3). Due to its geographical position, Mexico has historically served as a natural corridor for human migration, facilitating the movement of populations between North, Central, and South America. Understanding the country’s selection landscape is crucial to comprehend the genetic makeup of present-day American populations (4,5).

Recent advancements in next-generation sequencing technologies, along with the exponential increase of computational power, have provided the opportunity to identify patterns of natural selection in the human genome (6). In doing so, researchers have begun to understand not only the evolutionary history of populations but also the geographical distribution of genetic loci associated with phenotypic traits (7). While many studies have explored natural selection in populations around the world, few have focused on how it has shaped the genomes of present-day Mexican and MI populations (8–13).

Previous research has revealed distinctive selection patterns across different populations and regions within Mexico. Specifically, metabolic and immune-related genes under putative selection are consistently observed among MI populations. These signatures may have resulted from extended periods of metabolic or pathogenic pressures experienced throughout the population’s history, alongside exposure to novel pathogens introduced during La Conquista (8–13). Selection landscapes that are specific to physical endurance, stature, and metabolic efficiency have been identified in the Rarámuri, Triqui, and Seri Indigenous groups, respectively (9,10). However, these studies have based their inferences on SNP array and exome sequencing technologies, thus offering a limited perspective of selection across the whole genome (14). With a few exceptions (10,12,13), studies also limited their findings to ancestry-masked Mexican genomes with variable percentages of MI ancestry.

Here, we present the first comparative analysis of genetic adaptation in MI populations using whole genome data (WGS) with high Native American ancestry (>97%). Our study has enabled us to report selection signatures in previously unexplored regions of the genome, showing strong selective signals related to immune processes and providing further insight into possible genetic adaptations to pathogenic pressures. Moreover, we present nf-selection, a Nextflow pipeline for automated detection of recent selection using Population Branch Statistics (PBS) (15) and Integrated Haplotype Scores (iHS)(16). With nf-selection, we aim to streamline the identification of selection signatures on genome-wide data by simplifying the use of software needed for both tests. The pipeline is publicly available at GitHub: https://github.com/fernanda-miron/nf-selection.

## Results

### Data description

We analyzed 76 unrelated MI individuals from the **Metabolic Analysis in an Indigenous sample** (MAIS) cohort who were whole genome sequenced as part of a previous study (17). Individuals are representatives of 27 different ethnic groups in Mexico and show an average of 97.2% similarity to populations reported as Native American by ADMIXTURE (17). The individuals were divided into three previously defined geographic regions: Northern, Central, and Southern Mexico for this study (**Figure 1**) (17).

**Fig 1.**
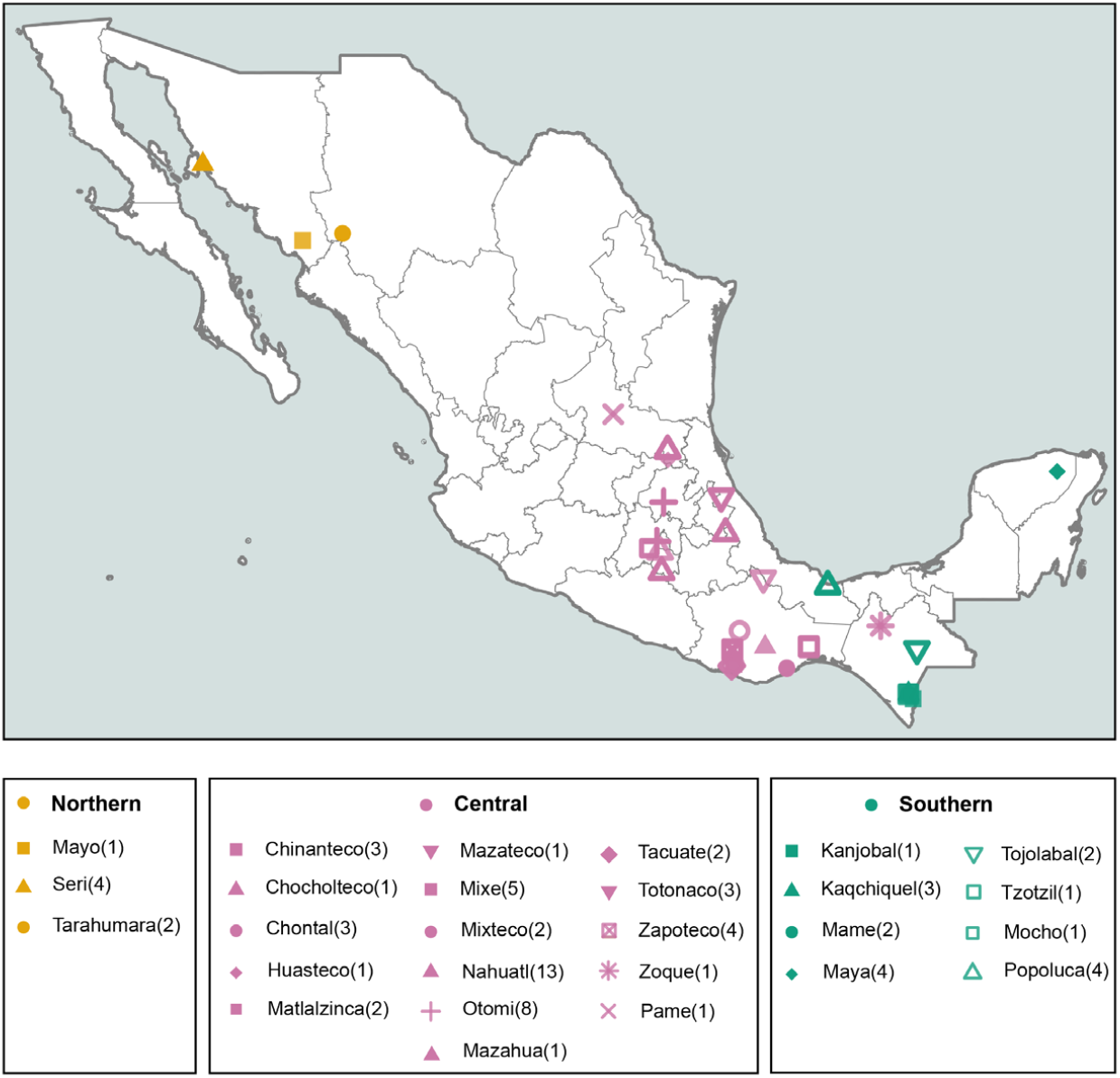
Approximate sampling locations of 76 unrelated MI individuals from the Metabolic Analysis in an Indigenous sample (MAIS) cohort. Individuals are representatives of 27 different ethnic groups in Mexico. Shapes denote the ethnic group, and colors denote their assigned geographic region. (N) indicates the number of individuals from each population. For selection analysis, samples were separated based on Northern (Yellow), Central (Pink), and Southern Regions (Green).

### Implementing a Selection Pipeline

To simplify the detection of selection signatures, we developed nf-selection, a Nextflow bioinformatics pipeline that computes two commonly used statistics: PBS and iHS. The PBS is a population differentiation-based test that identifies strong allele frequency differences between three populations by comparing pairwise F_ST_ values (15). The iHS is a linkage disequilibrium-based method that measures extended haplotype homozygosity at a given genomic region (16). For details on nf-selection implementation, review Methods. This pipeline is publicly available via GitHub (https://github.com/fernanda-miron/nf-selection) and includes instructions for installation and testing.

For each of our three study populations (Northern (N), Central (C), and Southern Regions (S)), we used nf-selection to calculate PBS for each autosomal variant using 33 unadmixed Peruvians (PEL) from the 1KGP as an ingroup and 50 Han Chinese (CHB) individuals from the 1KGP as an outgroup. iHS was also computed for each population cluster as part of the nf-selection pipeline. Genome-wide *P*-values were calculated separately for PBS using a distribution simulated under a demographic model specific to the Mexican Population (**Fig. S11**). Finally, we cross-referenced the top 1% of *P*-values for both statistics and focused on putative signals of selection detected by both methods to reduce our chances of reporting false positives (8).

### Selection Signatures Analysis

We found 2758 variants under putative selection for the Northern MI Populations, 1097 for the Central MI Populations, and 1459 for the Southern MI Populations (**Figure 2, Figs. S1-S6, Table S1-S3**). To prioritize putatively selected genomic regions for further analysis, we calculated the density of significant selection statistics in each region (i.e., the number of variants under selection normalized by gene length)(**Table S4-S6**). Genes with the largest value of selection density for each population are depicted in **Table 1** and **Figure 3**.

**Table 1.**
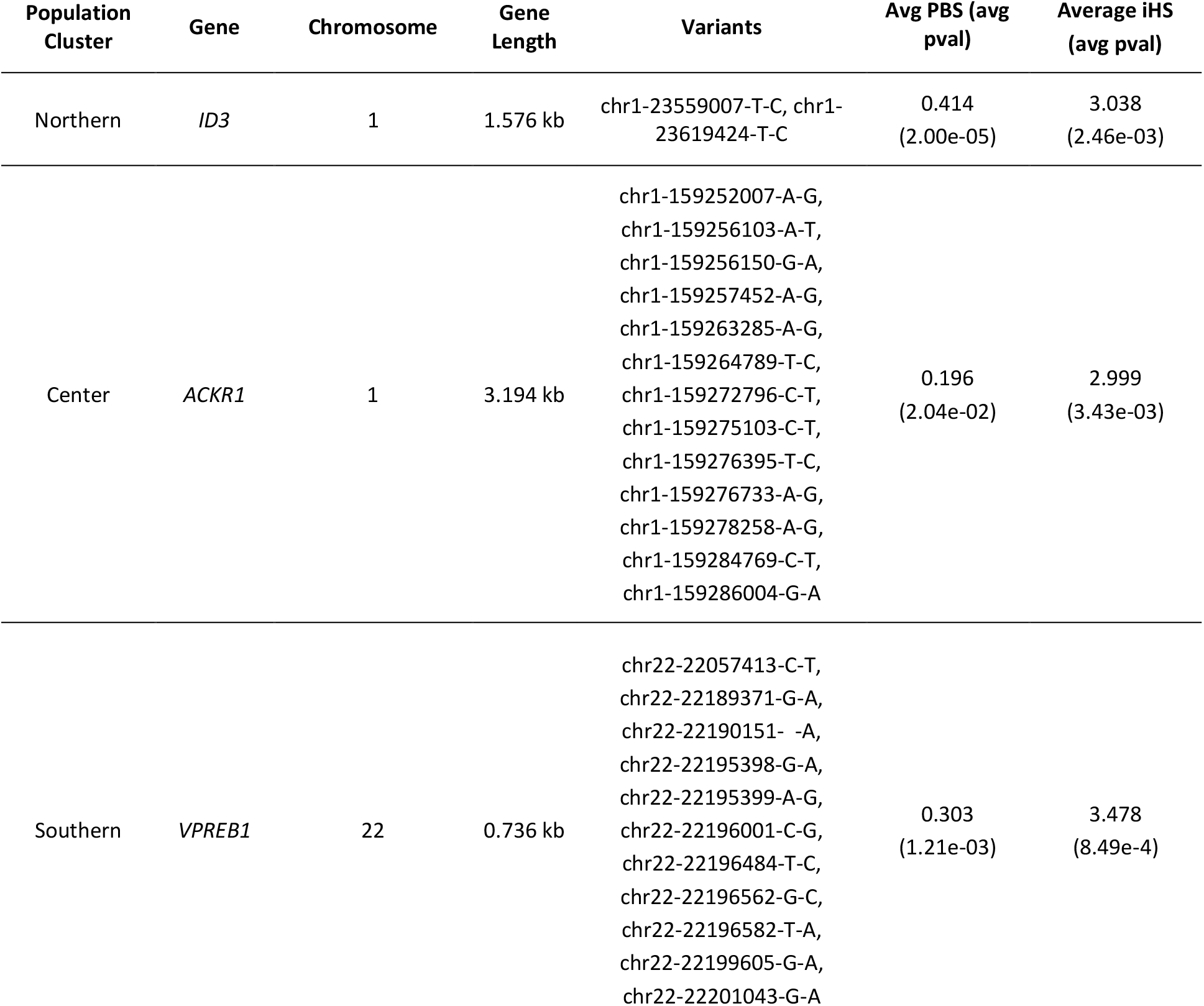
Top genes under selection in MI Populations based on Selection Density (Number of variants under selection normalized by gene length).

**Fig 2.**
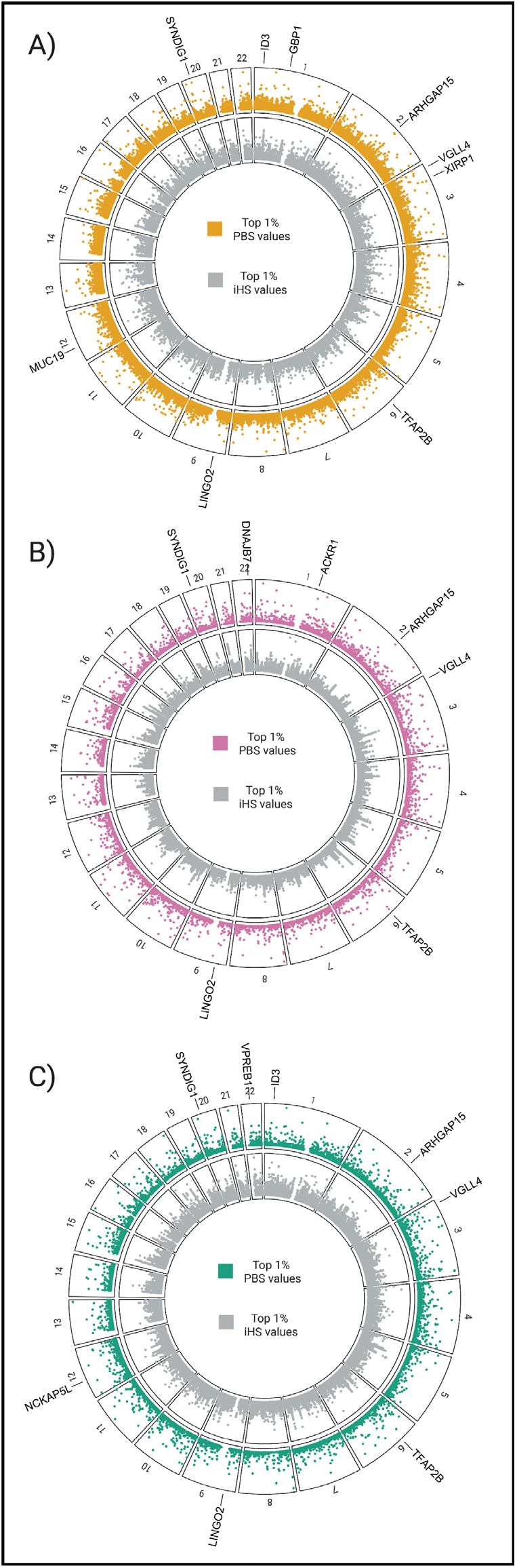
Significant adaptation signals in MI Populations. SNVs values under the top 1% distribution of PBS (Outer track) and iHS (Inner track) per autosome are plotted. Genes with the greatest selection density, with non-synonymous variants under putative selection, and shared between the three clusters are depicted. A) Northern MI Population B) Center MI Population C) Southern MI Population.

**Fig 3.**
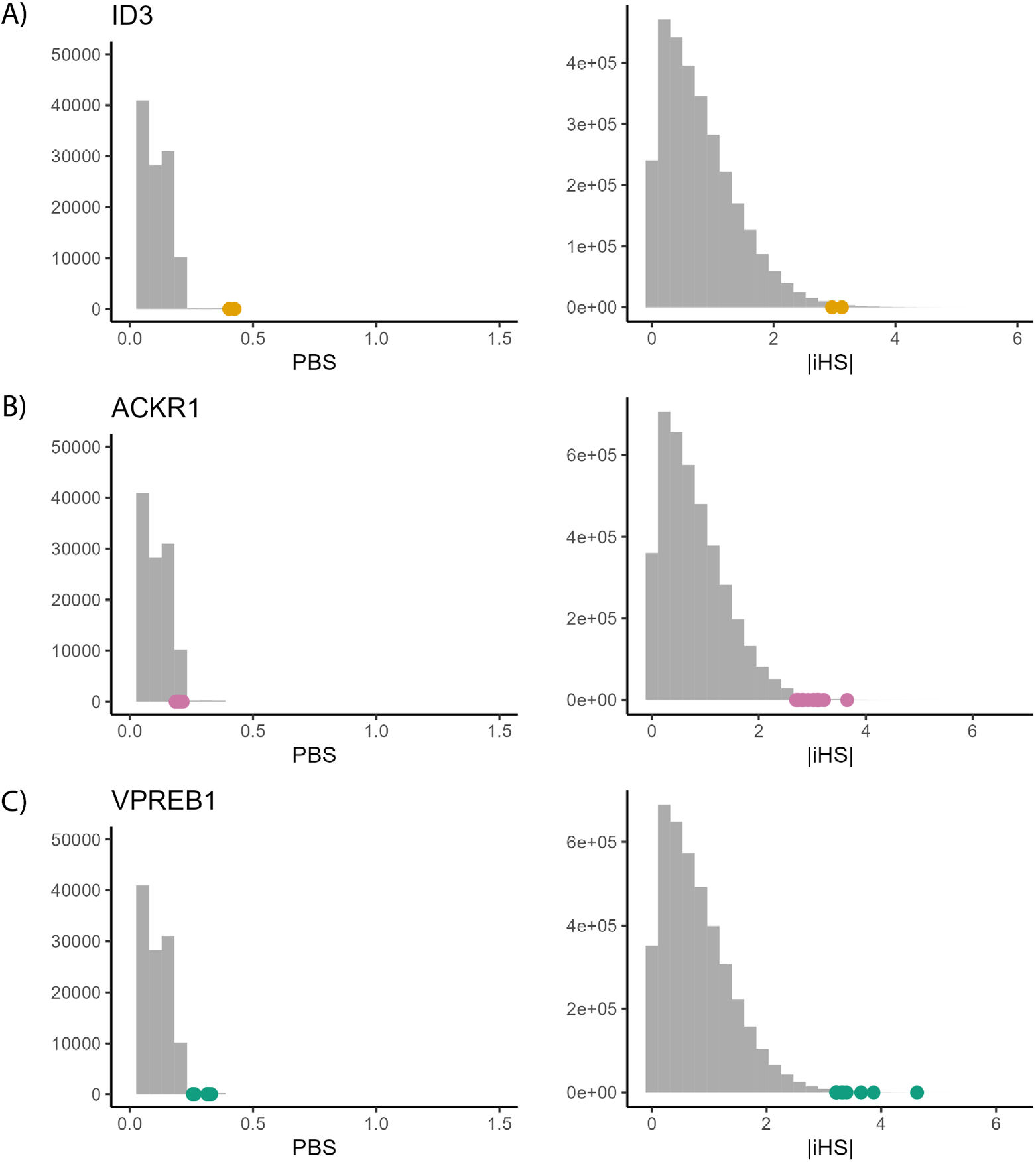
Genes with the strongest selection signals in MI Populations plotted against the simulated distribution of population branch statistics (PBS) and calculated distribution of iHS. Dots represent the values obtained for each SNP in the focal gene A) *ID3* (Northern MI Population), B) *ACKR1*(Center MI Population), C) *VPREB1* (Southern MI Population).

### Selection Signals in Regulatory Elements

Next, we looked for selection signals within enhancer and promoter regions using GeneHancer (v. 5.17), a novel database of human enhancers and their inferred target genes, in the framework of GeneCards (18). This analysis provides two types of results: one for variants per enhancer/promoter and one for variants per genomic feature. An enhancer/promoter element can regulate more than one genomic feature, and each genomic feature can be regulated by more than one enhancer/promoter. This was done independently for each region, and the results were compared across regions.

We find putatively selected variants in 382, 137, and 168 regulatory elements in the Northern, Center, and Southern Populations, respectively (**Fig. S7**). We calculated each regulatory element’s selection signal density (**Fig. S8**) and selected the top 10 hits per population (**Table S8**). The top 10 enhancers in every region are linked to non-coding elements, such as long non-coding RNAs (lncRNAs), microRNAs (miRNAs), and PIWI-interacting RNAs (piRNAs).

Next, we identified putatively selected variants in 21 variable regulatory elements shared between any two populations (**Table S7**). The Southern and Central Populations share three putatively selected SNPs in GH20J024425 (length 1999 bp) on chromosome 20, which regulate *SYNDIG1* and *GGTLC1*, two interferon-induced transmembrane proteins (20). We also identified eight putatively selected SNPs within five enhancers of the gene *MEI1*, known to be involved in meiosis and gamete generation (20).

Finally, we detected one enhancer, GH02J143479 (length 4,983 bp) (**Fig. S7**), containing putatively selected variants across the Northern, Center, and Southern populations, including the SNP chr2-143480377-T-C, which is shared across the three populations. This enhancer regulates the *ARHGAP15* and *GTDC1* genes and the non-coding elements *piR-44085-015, lnc-KYNU-7*, and *ENSG0000022865*. Interestingly, GH02J143479 exhibits one SNP under selection (chr2-143480377-T-C), consistently observed across all study populations, with the highest reported allele frequency (0.52) among the American Indigenous population in gnomAD v4.0 (19).

### Gene Annotation and Pathway Analysis

To further explore putative signals of selection identified in the study populations, we annotated significant variants as exonic, intergenic, intronic, etc., using Annovar (21) (**Table 2**). Consistent between the three study populations, most putative selective sites are located within intergenic regions. Additionally, among the variants on gene coding regions, intronic sites are overrepresented.

**Table 2.** Annovar annotation for variants under hypotheses of selection.

| Variant Type | Northern Variants | Central Variants | Southern Variants |
| --- | --- | --- | --- |
| UTR3 | 25 | 15 | 19 |
| UTR5 | 2 | 1 | 2 |
| downstream | 21 | 8 | 6 |
| exonic | 3 nonsynonymous SNVs, 3 synonymous SNVs, 1 unknown | 2 synonymous SNVs, 1 frameshift insertion | 2 nonsynonymous SNVs, 3 synonymous SNVs |
| intergenic | 1497 | 532 | 783 |
| intronic | 992 | 444 | 562 |
| ncRNA exonic | 15 | 6 | 6 |
| ncRNA intronic | 189 | 72 | 66 |
| upstream | 10 | 14 | 10 |
| Upstream/<br>downstream | 0 | 1 | 0 |

Since nonsynonymous mutations change protein sequences and are frequently the object of natural selection (22), we further focused our findings on non-synonymous variants found in our three study populations. For the Northern MI Populations, we found four non-synonymous variants in *GBP1, XIRP1, ID3*, and *MUC19*, respectively. A non-synonymous frameshift insertion in *DNAJB7* was found to be under selection in Central MI Populations, and two non-synonymous variants were identified in *ID3* and *NCKAP5L* for Southern MI Populations.

We conducted a KEGG pathway enrichment analysis using The Database for Annotation, Visualization, and Integrated Discovery (DAVID) (23) to study the pathways in which genes harboring signatures of selection may be overrepresented. However, after correcting for the false discovery rate (FDR < 0.05), we found no significant enrichment pathways for any of the three study populations.

Since adaptations to previous selective pressures may become maladaptations and contribute to disease susceptibility among populations, we also performed a Gene Disease Association scan using DAVID (**Tables S11-S13**). After FDR correction, we found significant associations for renal, psychiatric, metabolic, hematological, chemical dependency, and cardiovascular diseases in all study populations. Immune and neurological diseases were also significant in Northern and Southern MI Populations (**Fig. S10**).

### Shared Signals of Selection Between Populations

Next, we looked for genes with signals of selection among Northern, Center, and Southern Mexican populations by comparing statistically significant results from each analysis. Additionally, to increase the certainty of our analyses, we cross-referenced our results with previously reported signals of selection in MI Populations (8,9,12) (**Figure 4**).

**Figure 4.**
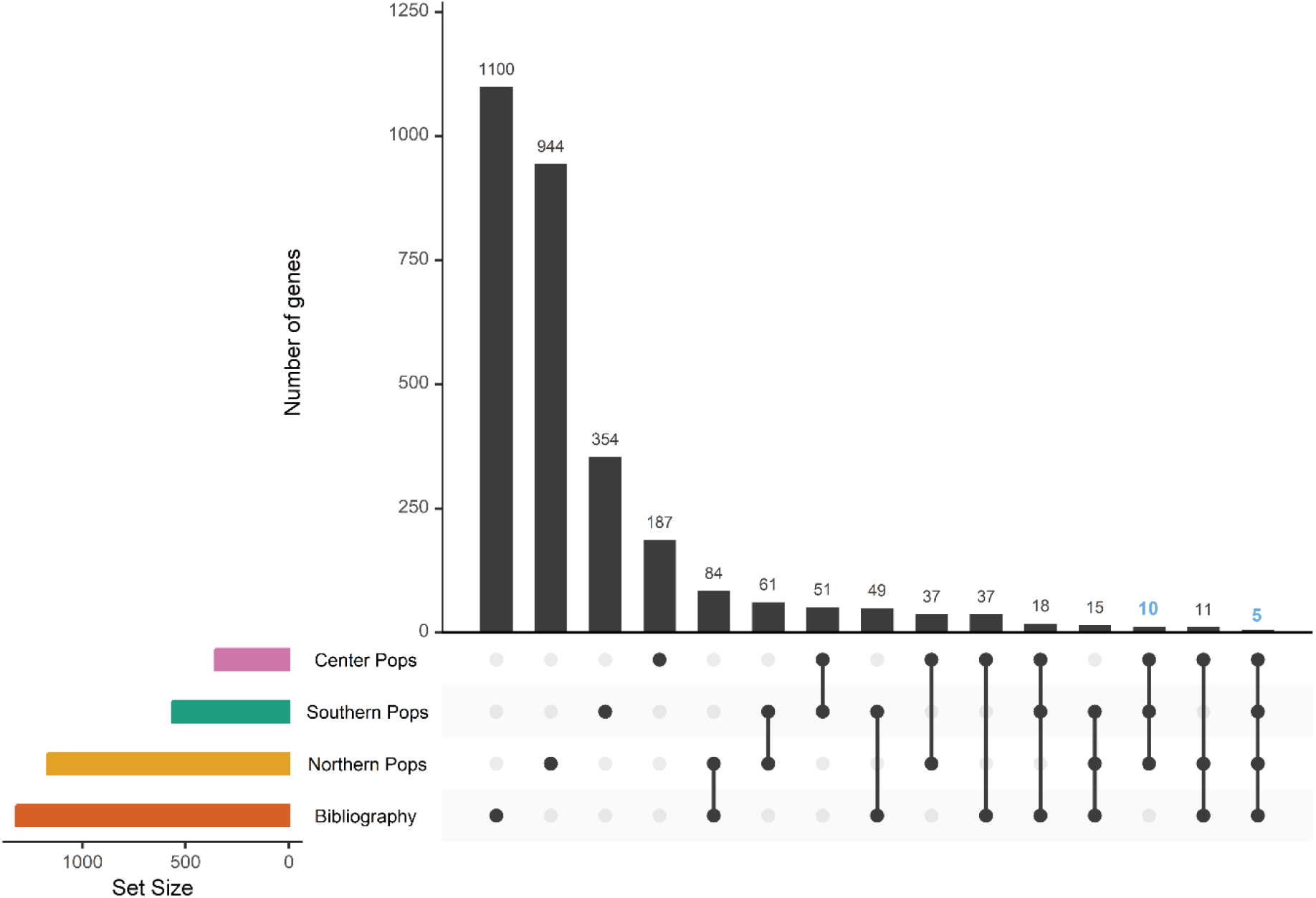
The intersection of Northern, Center, and Southern MI Populations, and previously reported genes (Bibliography) under putative selection on MI Populations. The bar plot (top) shows the number of genes per intersection. The matrix below indicates the intersection represented by each bar. The bar plots on the left show the total number of genes in every population and bibliography. The intersection between three study populations (10) and the intersection between three study populations and the bibliography (5) are depicted in blue.

We found ten genes with putative signatures of selection in all three populations(*CCSER1, KIF2B, LINC00430, LINC01721, LINC02546, LOC101928516, MIR8068, PKHD1, SLITRK6, WWOX*) that have not been previously reported in the literature, and five genes with signals of selection in all three populations that have been previously reported in the literature (*ARHGAP15, VGLL4, LINGO2, SYNDIG1, TFAP2B*). A brief overview and description of the function of each gene is provided in **Table 3**.

**Table 3.**
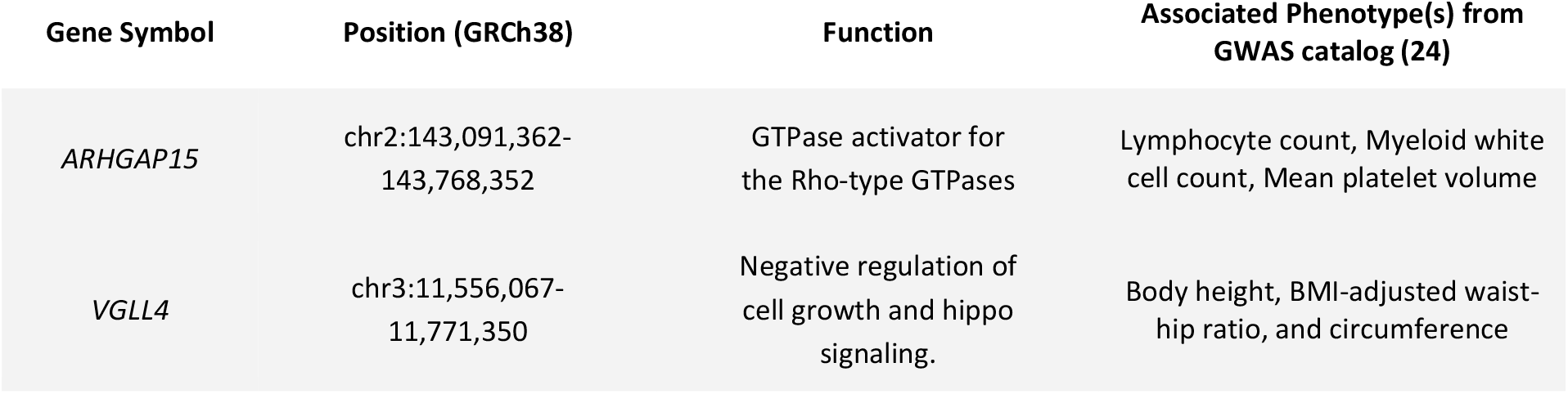
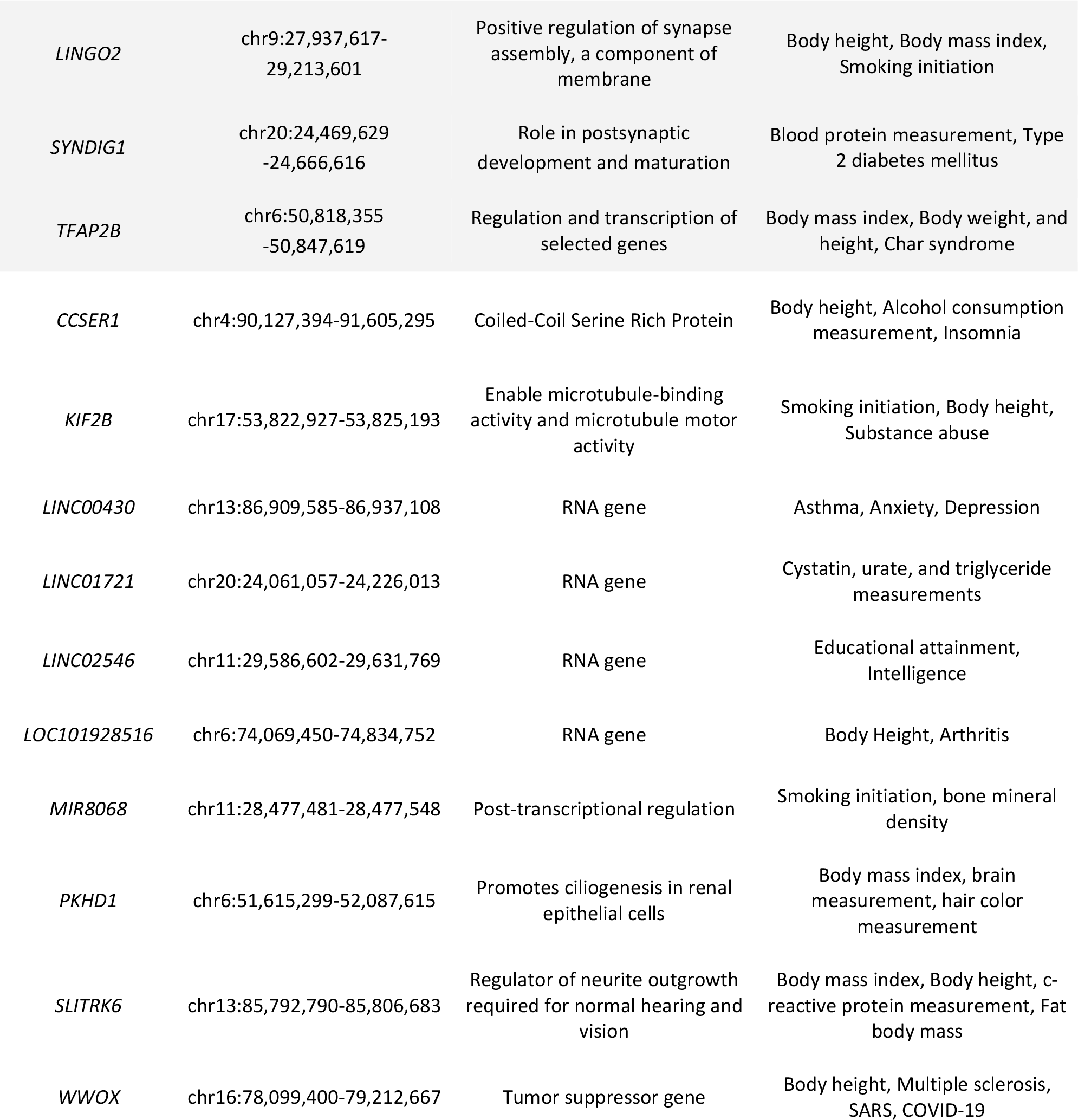
Genes with significant selection signatures across the three study populations. Genes previously documented in the bibliography are shaded in grey for reference.

## Discussion

The genetic structure observed in present-day MI populations reflects a combination of complex demographic and evolutionary events. The results of our study suggest that populations had collectively adapted to metabolic and immune pressures. However, specific selection landscapes were identified in different regions of Mexico.

### Northern MI Populations

The Northern MI Cluster shows non-synonymous variants putatively under selection on several key genes involved in immune defense (*ID3, GBP1, XIRP1*, and *MUC19*). Infectious diseases are among the strongest selective pressures shaping the human genome (29). Demographic events and cultural changes during recent human evolutionary history caused populations to be exposed to new and dangerous pathogens. Specifically, the European Colonization of the Americas represented a major epidemic challenge for MI Populations. After the arrival of the Spanish conquistador Hernán Cortés to what is today Mexico, dozens of epidemics swept through the country, leading to a population decline as high as 95% (30). This period also precipitated the introduction of several new pathogens, including the causal agents responsible for well-characterized diseases, such as measles, mumps, smallpox, and influenza (30).

Interestingly, both *GBP1* and *XIRP1* are involved in the immune response to *Salmonella* (26,27). *GBP1* (Guanylate Binding Protein 1) encodes a large GTPase of the dynamin superfamily involved in antimicrobial immunity and cell death. Specifically, *GBP1* promotes the detection of *Toxoplasma* DNA and the immune targeting of *Salmonella* (26). *XIRP1* (Xin Actin Binding Repeat Containing 1) is a protein-coding gene expressed mainly in fibroblasts and macrophages in response to cytokines and bacterial infections such as *Listeria, Shigella, and Salmonella* (27). A recent study by Vågene et al. (31) identified *Salmonella enterica* as a possible cause of the 1545–1550 “cocoliztli” epidemic in Southern Mexico. Historically “cocoliztli” epidemics were reported in the northern and central high valleys of Mexico, in addition to Southern Mexico (31), suggesting that this pathogen was a major selective pressure for Northern MI Populations during the colonial period.

Similarly, *MUC19* (Mucin 19) encodes a Mucin family protein that has been involved in the immune response to parasitic and viral infections (28). The *MUC19* gene that we identify as putatively under selection in the Northern cluster, has been previously identified under selection in both populations from central Mexico (8) and individuals of Mexican heritage living in Los Angeles (MXL) from the 1000 Genomes Project (32). In a recent article, Villanea et al. (33) described the presence of a *Denisovan-like MUC19* haplotype at high frequencies in admixed Latin American individuals. Their results suggest that this haplotype served as raw genetic material for positive selection in American populations.

### Central MI Populations

The Central MI Cluster shows variants under selection in the *ACKR1* and *DNAJB7* genes. *ACKR1* (Atypical Chemokine Receptor 1), formerly referred to as DARC (Duffy antigen receptor for chemokines), encodes a glycoprotein expressing the Duffy blood group antigens. The Duffy protein acts as a receptor for the malaria parasites *Plasmodium vivax* and *Plasmodium knowlesi* (34). Malaria, a global infectious disease caused by *Plasmodium spp*. and spread to humans by some types of mosquitoes, has been one of the most important selective pressures on the human lineage (35). *ACKR1* has previously been reported as a target of directional positive selection in sub-Saharan African populations with resistance to erythrocyte invasion by *Plasmodium vivax* (36). Interestingly, our Center MI cluster shows several variants under putative selection in *ACKR1*.

In Mexico, Malaria continues to pose a significant risk for indigenous peoples living in Chiapas and Oaxaca, with *Plasmodium vivax* being the dominant *Plasmodium* all over the country (37). Based on the World Health Organization (WHO) Malaria Report, there have been a cumulative total of 8,157 indigenous malaria cases in Mexico recorded from 2010 to 2022 (38). How and when *P. vivax* arrived in the Americas remains highly controversial, however, most theories propose pre- and post-Colonial exposures with Asian and European contributions to the genetic diversity of the parasite (39,40). Our results from the Central study cluster suggest that a malaria may be driving variation at *ACKR1*. Nevertheless, present-day American Indigenous Populations do not show evidence of protective phenotypes for malaria — Sickle-cell trait, glucose-6-phosphate dehydrogenase (G6PD) deficiency, and *ACKR1* negative expression — so further work is required to determine to functional impact, if any of the putatively selected allele (39).

*DNAJB7* (DnaJ Heat Shock Protein Family (Hsp40) Member B7) is an intronless gene that belongs to the DNAJ/HSP40 family of proteins, which regulate chaperone activity (18). Heat shock proteins (Hsps) play a vital role in cell homeostasis under both physiological and stressful conditions (41,42). Particularly, the DnaJ Heat Shock Protein Family, or Hsp40, constitutes the largest and most diverse subgroup of Hsp families, playing fundamental roles in neurodegeneration, tumorigenesis, glucose homeostasis, and spermatogenesis (43,44). *DNAJB7*, a gene with a non-synonymous mutation under selection in the Central MI Populations, is a member of the DnaJ family that has been related to the pathogenesis of insulin resistance and Type 2 Diabetes Mellitus (T2D) (43).

Genes related to metabolism and, more specifically, T2D, have previously been reported under putative selection in MI Populations as a result of adaptation to periodic starvation experienced throughout history (12,45). The rapid change in diet and behavior in the Contemporary MI populations against a genotype that cannot adapt rapidly creates an evolutionary mismatch expressed as metabolic diseases (46). Additionally, *DNAJB7* is highly expressed in the testis, suggesting a possible role in male fertility (47). Considering these observations, selection under *DNAJB7* may result from metabolic or reproductive selective pressures.

### Southern MI Populations

The Southern MI Cluster shows variants under putative selection in the *VPREB1, NCKAP5L*, and *ID3* genes. *VPREB1* (V-Set Pre-B Cell Surrogate Light Chain 1) encodes a protein belonging to the immunoglobulin superfamily, expressed at the early stages of immune B cell development (18). *NCKAP5L* (NCK Associated Protein 5 Like) is a gene that regulates microtubule organization, stabilization, growth, and bunding formation (18). The gene *ID3* (Inhibitor of DNA binding 3) encodes a transcription factor of the helix-loop-helix family reported to be involved in the differentiation and growth of a variety of cell types (18). The coding product of *ID3* controls the formation of effector and memory CD8+ populations, a critical process for adaptive immunity (25). As in Northern and Center MI Populations, pathogenic environments seem to be a major selective pressure for Southern MI Populations. To the best of our knowledge, *VPREB1* and *ID3* are two genes heavily involved in different immune-related pathways that do not have reports of pathogenic specificity.

On the other hand, *NCKAP5L* is a gene that has been associated with *Spherocytosis* Type 2 (SPH2)(OMIM #616649) (18), a hereditary disease characterized by the presence of spherical-shaped erythrocytes on the peripheral blood smear and a possible protective phenotype against *Plasmodium falciparum* (48,49). The deadliest of the human malaria parasites, *P. falciparum*, presumably entered the Americas after European contact as a result of the trans-Atlantic slave trade (39). During the Colonial period in Mexico, the slave trade was primarily through the southeastern ports of Veracruz and Campeche seaports (50) leading to sugar plantations and other extractive industries in the Yucatan and beyond (51). This history may suggest that Southern MI Populations have had extended exposure and potentially selective pressures from *P. falciparum* infection than other regions of Mexico.

### Shared Selection Signatures between MI Populations

When comparing signals of selection across our study regions, we identified 10 (*CCSER1, KIF2B, LINC00430, LINC01721, LINC02546, LOC101928516, MIR8068, PKHD1, SLITRK6, WWOX*) genes with selection signatures that were present in all our three study populations, and five genes that, besides being present, had previous reports of selection on MI Populations (*ARHGAP15, VGLL4, LINGO2, SYNDIG1, TFAP2B*). Our results showed an overrepresentation of genes involved in metabolic and immune pathways.

*ARHGAP15* is a gene that encodes an RHO GTPase-activating protein highly expressed in immune reservoir tissues (20). Particularly, neutrophils in mice with an *ARHGAP15* knock-out showed an increase in bone marrow retention (52). Under healthy conditions, most neutrophils are located in the bone marrow (retention). However, during infection, they are rapidly released into the bloodstream to combat pathogens (egress). For both pathways to work, common intracellular signaling elements, such as (GTPases) of the Rho family, are essential. This information confirms the role of *ARHGAP15* in immune cell mobilization. Our results on regulatory elements also show some variants under selection on enhancer elements connected to *ARHGAP15*. Moreover, selection in this gene has previously been reported in MI Populations (8,9,12 & Figure 5).

Genome-wide association studies (GWASs) showed that different SNPs in *LINGO2* were associated with obesity, T2D, and gestational diabetes mellitus risk (53). Moreover, *LINGO2* has been related to immunity against helminth infection (54,55). *VGLL4, TFAP2B*, and *WWOX* have been implicated in increased Body Mass Index, Insulin Resistance, Lipid and Triglycerides levels, and increased risk for Type 2 Diabetes (23,56,57). Since metabolic phenotypes under selection result from physiological adaptation to the available food supplies, we hypothesize that metabolic adaptation may have occurred either before the divergence of our three clusters. Signals of selection in *LINGO2* and *ARHGAP15* may also be due to selective pressures from the pathogens brought to Mexico by the conquistadors.

### Selection in Regulatory Elements

Using the additional data afforded us by WGS data, we investigated signals of selection within enhancer and promoter sequences, to better understand the selective landscape of regulatory elements in MI populations. Our results show that most of the altered enhancer elements of interest are connected to non-coding features, with notable exceptions like *ARHGAP15, GTDC1, SYNDIG1, GGTLC1, MEI1, ADSL, TOB2*, and *NALF1*.

*GTDC1* is a glycosyltransferase-like encoding gene highly expressed in the brain, fat, and testis. This gene, predicted to enable glycosyltransferase activity, has been associated with different neurodevelopmental disorders (58), and *Mycobacterium tuberculosis* infection resistance (59). *SYNDIG1* encodes an interferon-induced transmembrane protein, highly expressed in the brain and muscle (20). *GGTLC1* encodes a gamma-glutamyl transpeptidase interferon-induced transmembrane, highly expressed in the lung and testis (18). *MEI1*, targeted by five different enhancers with SNPs under putative selection, is a protein-coding gene predicted to be involved in gamete generation, meiotic spindle organization, and meiotic telomere clustering. Polymorphic alleles of *MEI1* have been linked to human azoospermia, a medical condition characterized by the absence of sperm in the ejaculate due to meiotic arrest (60). Furthermore, this gene was previously identified as belonging to the Human Phenotype Ontology (61) category “Female infertility”. These associations suggest putative selection on regulatory regions associated with reproductive phenotypes.

## Conclusions

This analysis represents the first whole-genome selection scan performed on MI Populations. Altogether, our study found signals of selection in metabolic and immune-related genes. However, specific selection landscapes were identified in different regions of Mexico. For the Northern MI cluster, we found evidence of selection on genes involved in the specific immune response to Salmonella and the unspecific immune response to different infectious diseases. Both Central and Southern clusters exhibit a selection landscape putatively shaped by different malaria strains and other pathogens.

To aid in this and future studies of natural selection, we also presented nf-selection, a pipeline that allows the computing of PBS and iHS to detect selection signals in diploid organisms. The developed pipeline offers a friendly shortcut to the reproducible and organized application of the bioinformatics code required to apply both statistics. Particularly, the use of Nextflow as a workflow orchestration engine in the pipelines provides the possibility of parallelization to speed up analyses. We showed that nf-selection represents a practical tool for the detection of selection signals, and we expect that it will prove useful for the research community.

We recognize several limitations that were encountered during this investigation. Firstly, the number of whole genomes available for analysis is limited for multiple ethnic Indigenous groups. This may restrict the representation of the 27 Mexican populations in the findings described. Additionally, the significant genetic drift exhibited by the Seri and Tarahumara populations (17) of the Northern MI group poses a challenge in establishing robust conclusions for the Northern cluster. This diversity necessitates careful consideration when discussing the region-specific results. Secondly, while several signals of selection have been identified on immune genes that may be associated with the recent colonial history of Mexico, future simulation work is needed to estimate the timing and strength of selection on these loci. Finally, as with all studies of genome-wide selection scans, our results only provide the first step to understanding the phenotype resulting in selective pressure. Future work with functional genomics and deep phenotyping will be required to fully understand the biological consequences of putatively selected variants. Despite these limitations, this research provides valuable insight into the adaptation processes that MI populations experienced in the recent past, paving the way for further WGS selection analysis.

## Code Availability

The Nextflow pipeline to reproduce this work is publicly available at: https://github.com/fernanda-miron/nf-selection

## Data Availability

New genomic data was not produced. Ethics statement and access to data can be found at the original data source (17).

## Methods

### Samples

Samples in this study were previously reported by Aguilar-Ordoñez I. et al. (2022)(17). In brief, 76 individuals from the MAIS cohort were whole-genome sequenced. Individuals are representatives of 27 different ethnic groups in Mexico and show an average of 97.22% similarity to populations reported as Native American by ADMIXTURE (17).

### QC and data treatment

Whole genome data in VCF format was filtered with <u>bcftools v1.8</u> to obtain data from autosomal chromosomes. The file was merged with whole genome data from 33 Peruvian individuals without any evidence of recent European or African genetic ancestry selected from the 1000 Genomes Project (1KGP) and with 50 Han Chinese individuals from the 1KGP using <u>bcftools v1.8.</u> The merged VCF was filtered to keep positions with at most 0.8% of missing data and MAF=0.05. Using these criteria, the final dataset contained 4,624,975 variants.

### Selection pipeline

To simplify the detection of selection signatures, we developed nf-selection, a Nextflow pipeline that detects candidate genomic regions and genes under selection by computing two commonly used statistics: PBS and iHS. The analysis starts with haplotype phasing with SHAPEIT 4.2 (62), followed by ancestral allele annotation using fasta ancestral files and the JavaScript vcfancestralalleles.jar from Jvarkit (https://github.com/lindenb/jvarkit). PBS calculation is implemented in unfiltered ancestral/derived variants with an in-house R script using Wright Fst values obtained from vcftools (63). The calculation of iHS is carried out using the R package rehh (65), using filtered ancestral/derived variants for iHS. Genome-wide P-values are calculated separately for PBS using a distribution simulated under a demographic model specific to the population of interest. Genome-wide P-values for iHS are retrieved from the rehh output. To reduce false positives, we follow the cross-reference approach of Reynolds et al. 2019 (8), cross-referencing between the top 1% of PBS and iHS. Cross-referenced SNPs are annotated using ANNOVAR. These results can be used for downstream analyses of gene curation, gene-set over-representation, gene networks, etc.

### Demographic model

A demographic model was modified from Reynolds et al. (8) to calculate expected distributions of PBS under neutral demographic processes for our three study regions using <u>fastsimcoal2</u> (version fsc27.09). Briefly, authors retrieved previously published estimates for the timing of the Out-of-Africa bottleneck and peopling of the Americas events (Timing of the bottleneck coinciding with the peopling of the Americas, time of the population divergence between PEL and the study population, timing of a possible bottleneck for the study population) (8,9)(**Fig. S11**). Joint site frequency spectrum of the CHB, PEL, and the study populations were calculated using <u>easySFS</u> (https://github.com/isaacovercast/easySFS). We included the divergence time between the PEL and our study population as an open parameter ranging between 480 and 520 generations. Additionally, the timing and severity of the most recent bottleneck were left as an open parameter in the model, as genetic and anthropological evidence for many Indigenous populations remains uncharacterized. Unknown parameters for the MI Populations were estimated by running the model 100 times with 1,000,000 iterations per run. The best likelihood run was chosen and used to simulate 1,000,000 sites across 22 chromosomes for 100 CHB, 66 PEL, and 152 MI individuals. The simulation was done 100 times, and variant sites from one randomly selected simulation were chosen to calculate PBS. Values of PBS were used to form a distribution for comparison with the observed empirical values.

### Whole genome scan for selection signatures

Using our pipeline, we computed PBS and iHS for the three Mexican regions. We calculated the PBS for each autosomal variant in each population using 33 unadmixed Peruvians from the 1000GP as an ingroup and 50 Han Chinese individuals from the 1000GP as an outgroup. The pipeline was run with default settings. Moreover, *P*-values for PBS for each region were calculated using a distribution of PBS values under a neutral demographic model. P-values for iHS were extracted from the rehh output. The top 1% of P-values for PBS and iHS were identified and cross-referenced to keep only entries that were present in both statistics. This cross-referenced approach has been used previously to avoid false positive results (8).

### Detection of variants in enhancers

We detected which SNPs are located in an Enhancer or Promoter element for each regional selection signal dataset using a pipeline that cross-references variant positions with the GeneHancer database (v5.17, downloaded Sep 01, 2023). The pipeline can be downloaded from https://github.com/Iaguilaror/nf-vcf2genehancer. In brief, using R scripts, the GeneHancer v5.17 GFF file is converted to a pseudo-BED format to keep the connected genes information, then the query VCF file is cross-referenced with the pseudo-BED using bedtools intersect (v 2.31.0); finally two summaries of variation are created: one for variants per GeneHancer element, and one for variants per genomic feature (a GeneHancer element can regulate more than one genomic feature, and each genomic feature can be regulated by more than one GeneHancer element). The process was run independently for the north, central, and south datasets. The comparison of regulatory variation between geographic regions was performed with an in-house R script.

## Acknowledgments

We thank the communities involved in this research and the participants who provided samples for analysis. We thank Olvera-Acosta Ricardo Benjamin and Orozco-Flores Diego from the Expression Analysis Subdirectorate at the National Institute of Genomic Medicine (INMEGEN) for helping organize the data and Denise García Castro, Andrea Arriola-Gamboa, and José de Jesús Mares Guerra for curating the data. This work was partially performed at cluster INMEGEN, which receives technical support from Gómez-Romero Laura.

