## Supplemental Figures 1-11 for "Whole genome sequencing of 76 Mexican Indigenous reveals recent selection signatures linked to pathogens and diet adaptation"

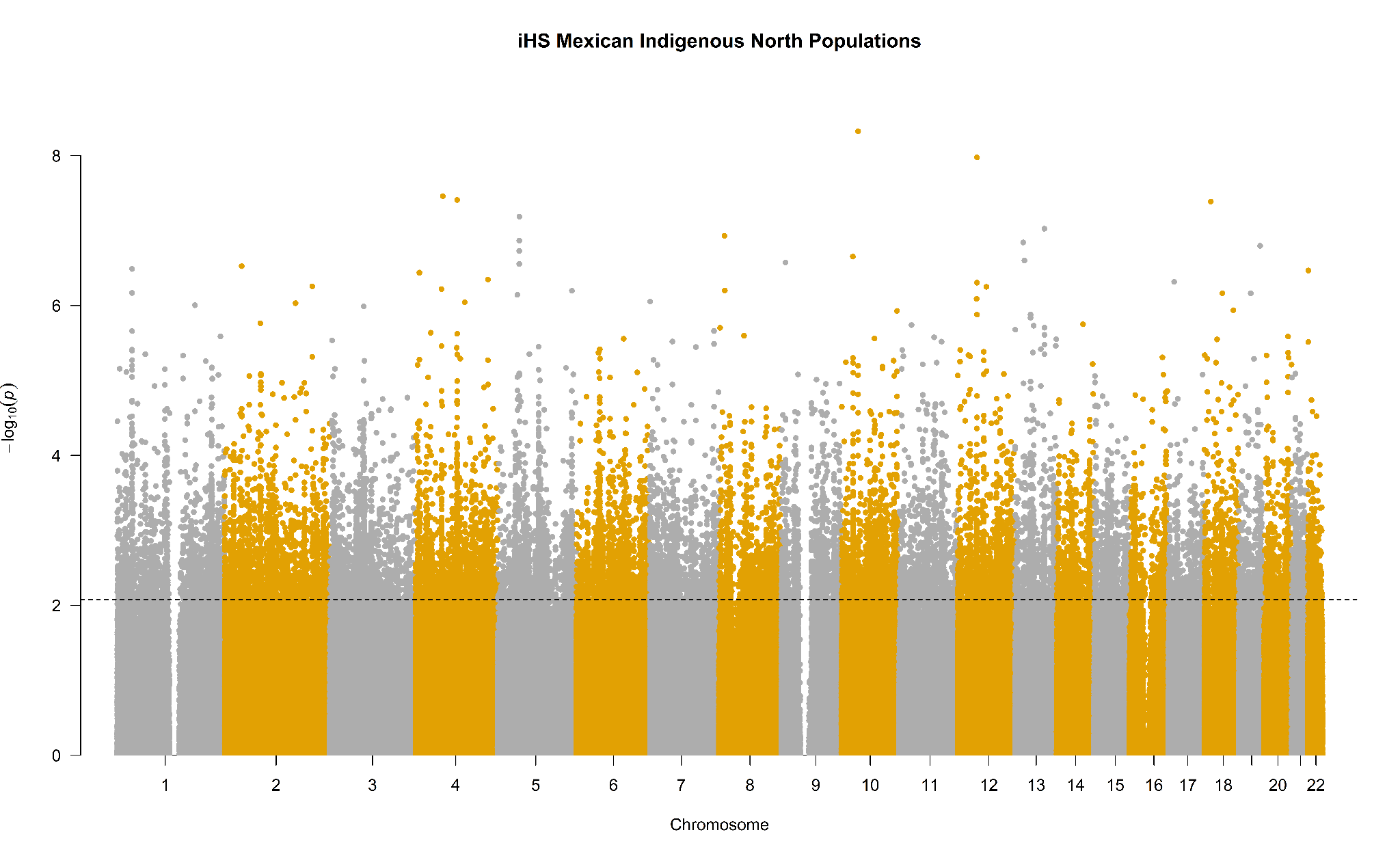

**Figure S1.** Manhattan plot of the iHS p-values for the Northern Mexican Indigenous Populations. The dashed line represents the 1% cutoff.

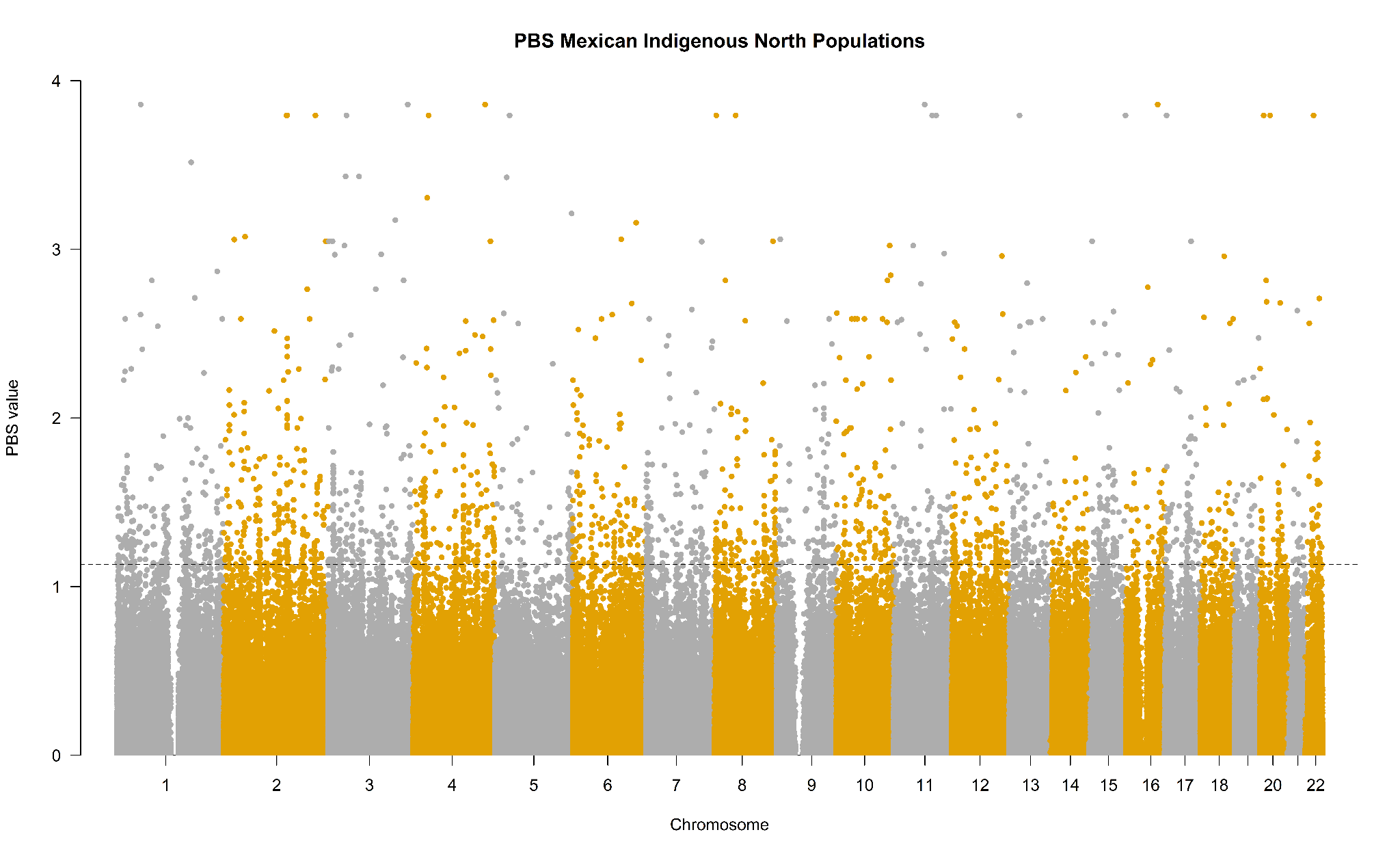

**Figure S2.** Manhattan plot of the PBS values for the Northern Mexican Indigenous Populations. The dashed line represents the 1% cutoff.

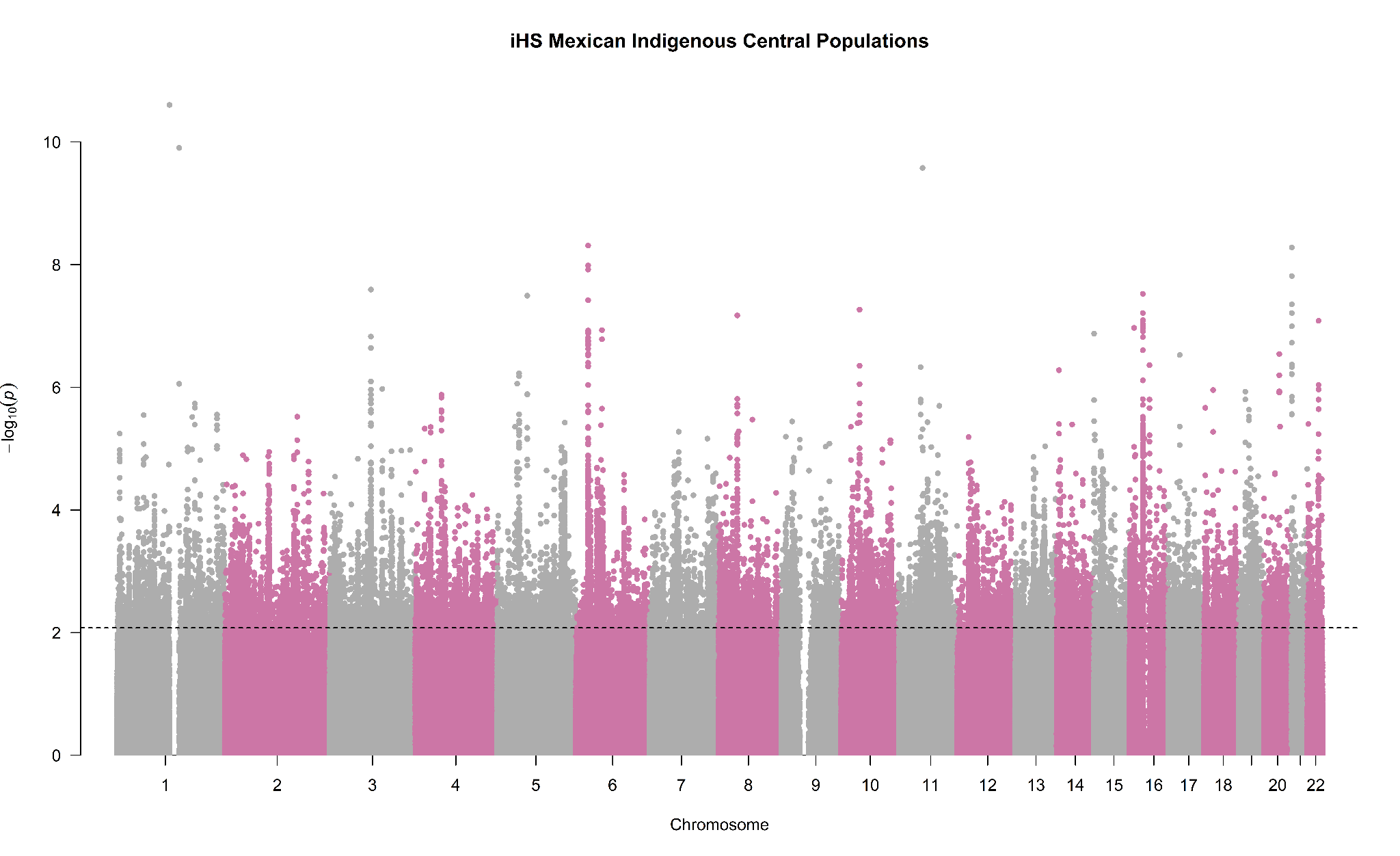

**Figure S3.** Manhattan plot of the iHS p-values for the Central Mexican Indigenous Populations. The dashed line represents the 1% cutoff.

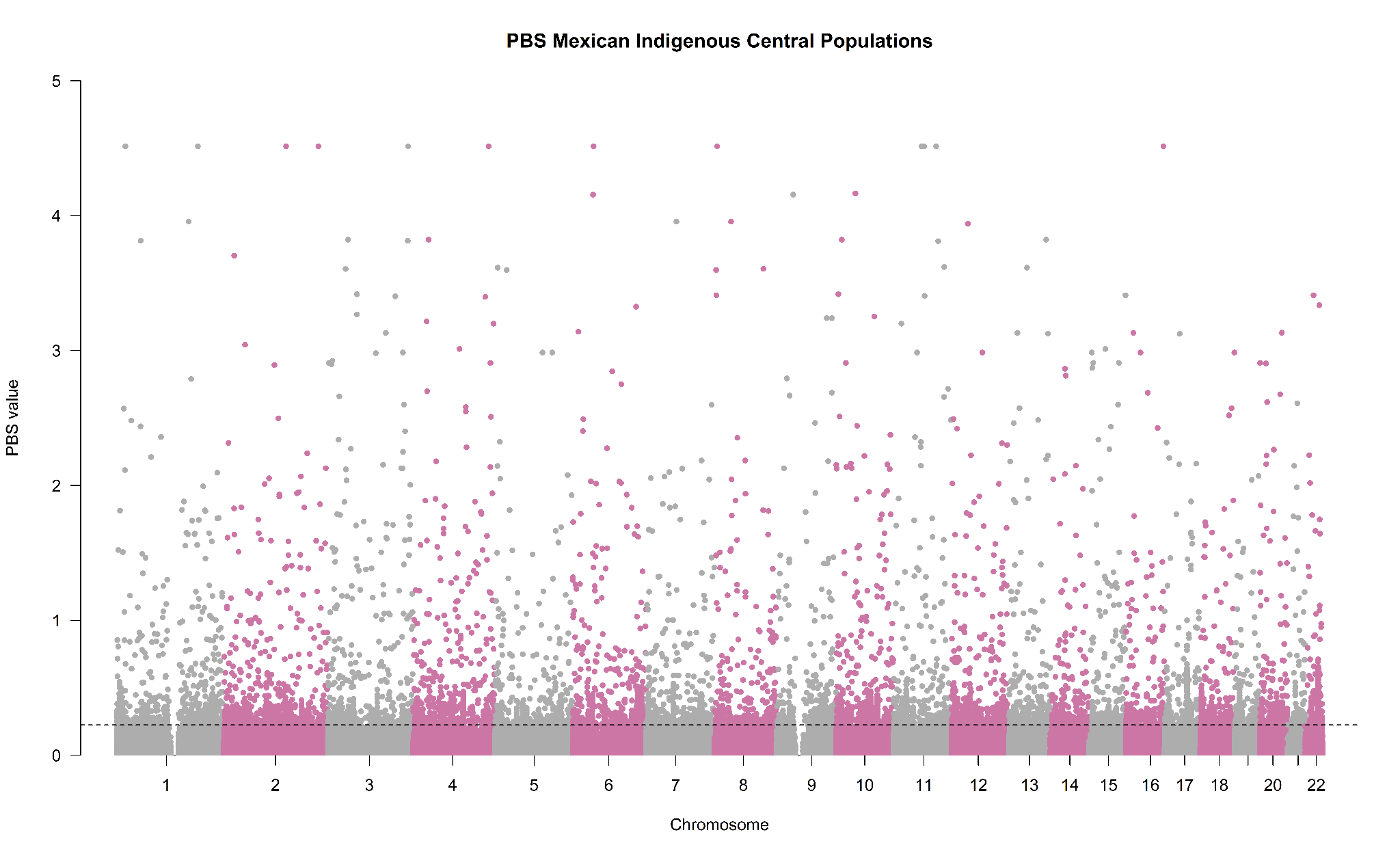

**Figure S4.** Manhattan plot of the PBS values for the Central Mexican Indigenous Populations. The dashed line represents the 1% cutoff.

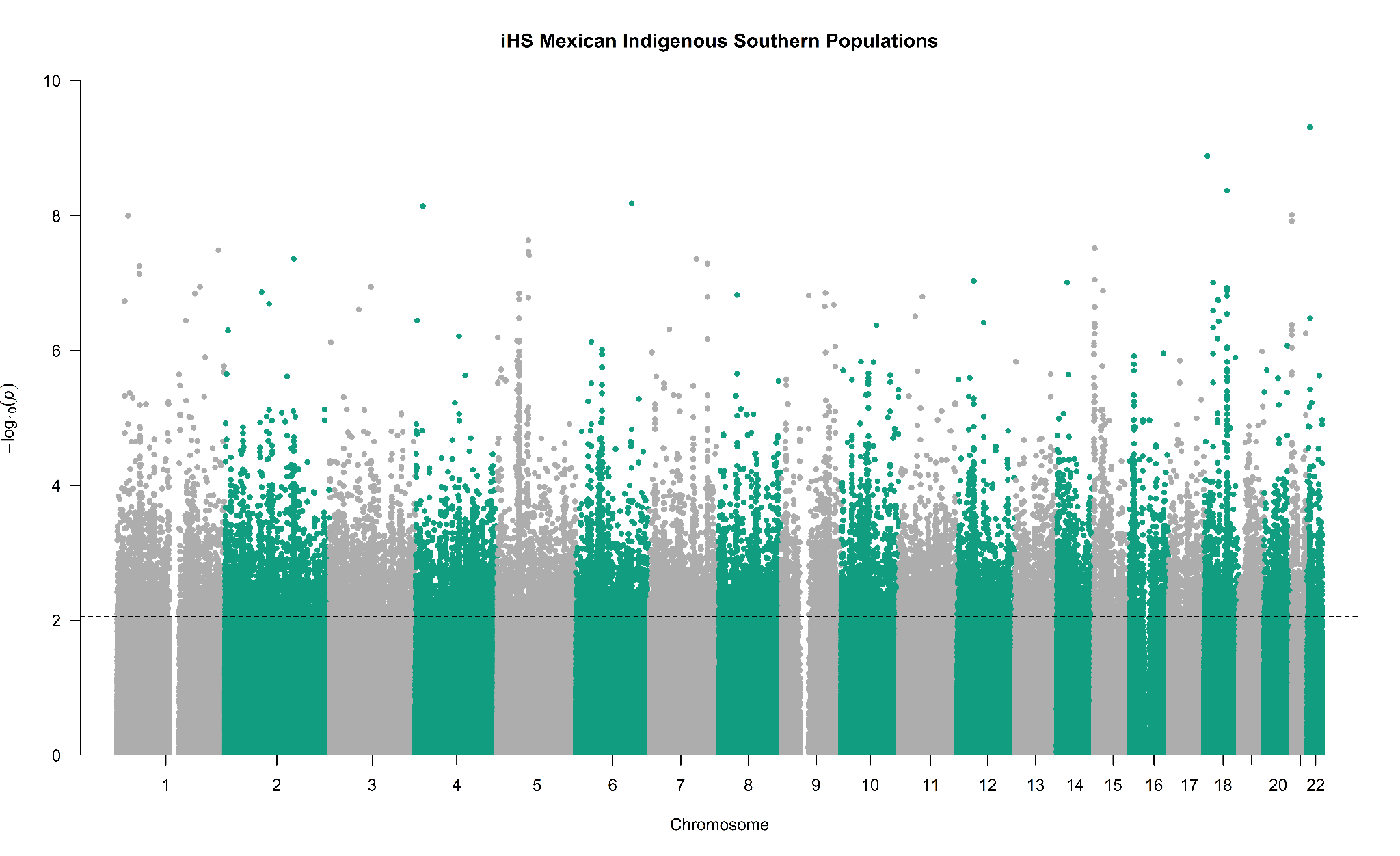

**Figure S5.** Manhattan plot of the iHS p-values for the Southern Mexican Indigenous Populations. The dashed line represents the 1% cutoff.

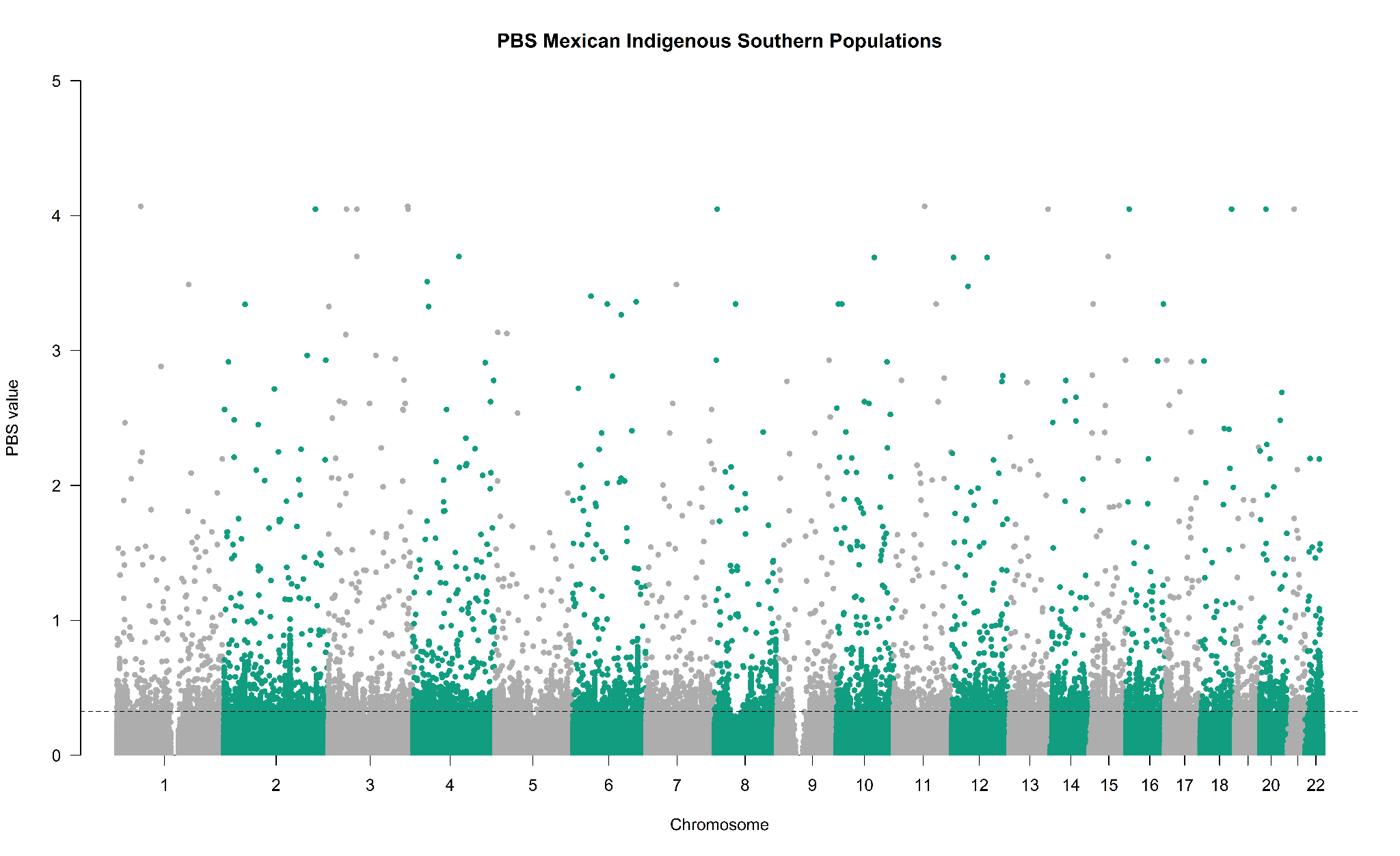

**Figure S6.** Manhattan plot of the PBS values for the Southern Mexican Indigenous Populations. The dashed line represents the 1% cutoff.

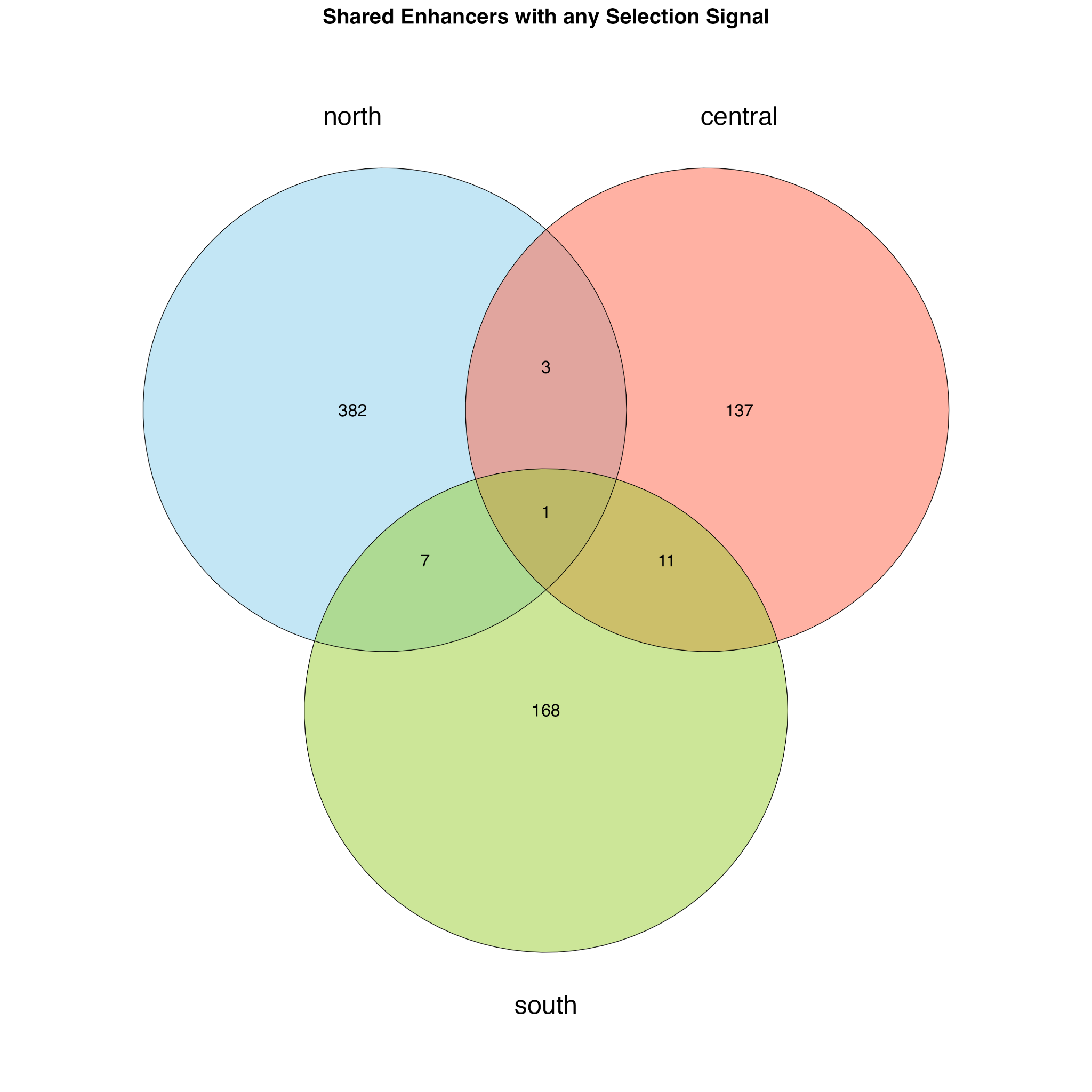

**Figure S7. Shared regulatory elements with selection signals.** Some enhancers are altered in more than one of the geographic regions, while some are exclusive to a region.

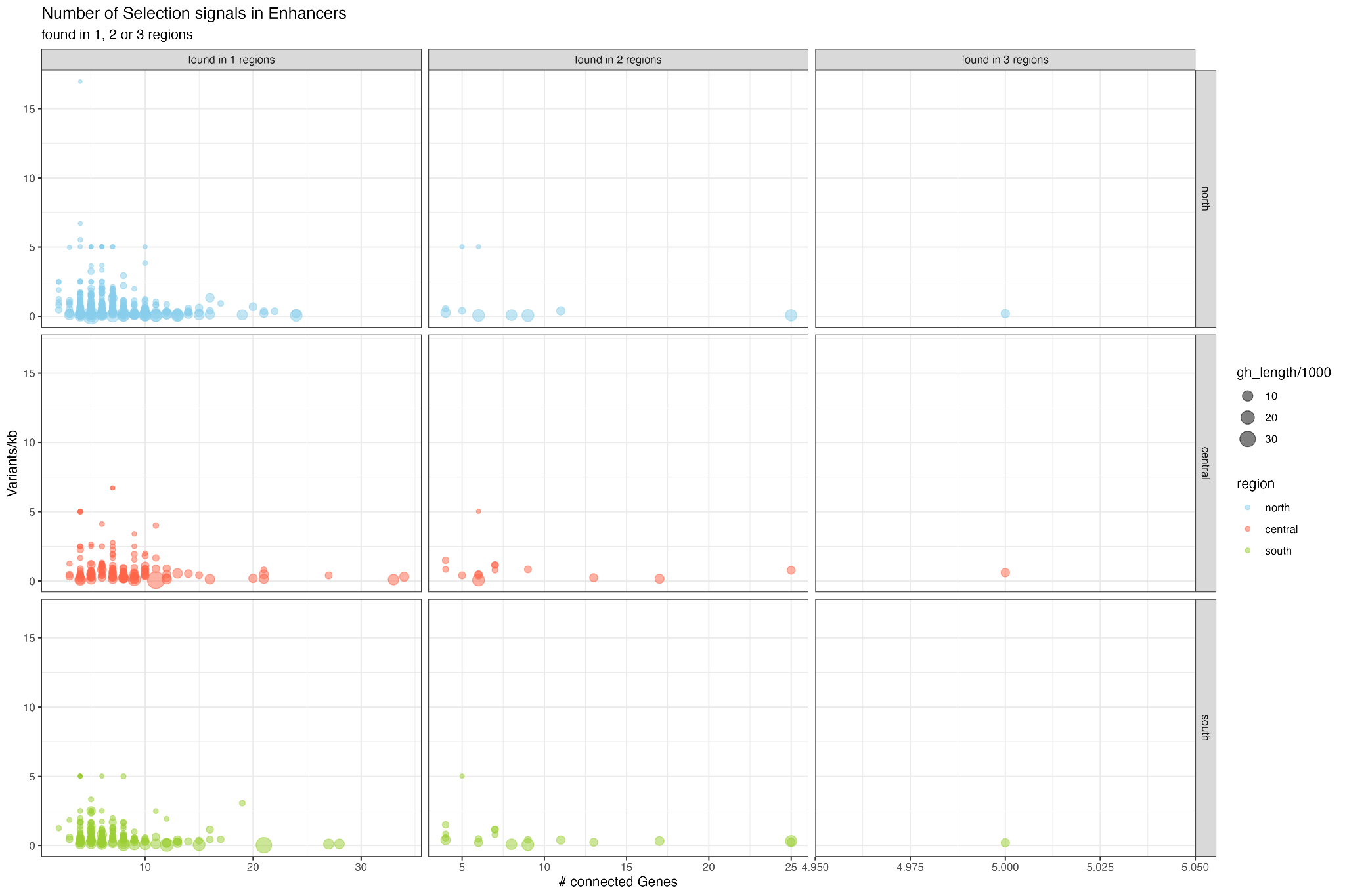

**Figure S8. Selection signal density by region.** We measured the number of selection signal SNPs divided by the length of the regulatory element (y-axis). We found that selection signals shared by 2 or 3 geographic regions show lower selection signal density.

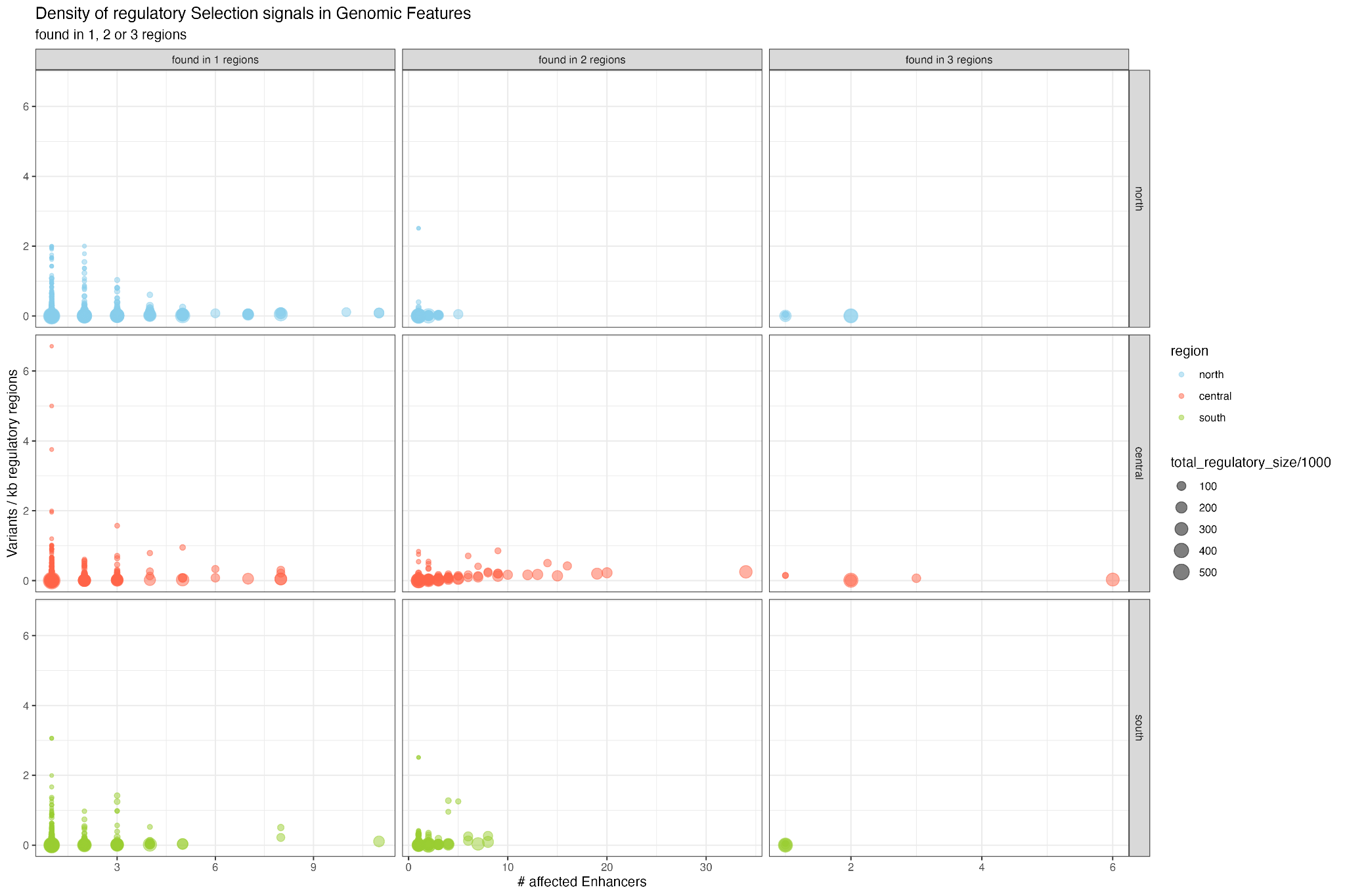

**Figure S9**. Regulatory selection signal density in genomic features by region. We measured the number of selection signal SNPs divided by the summarized length of all the regulatory elements for a given feature (y axis) and found that altered regulatory elements shared by 2 or 3 geographic regions show lower selection signal density.

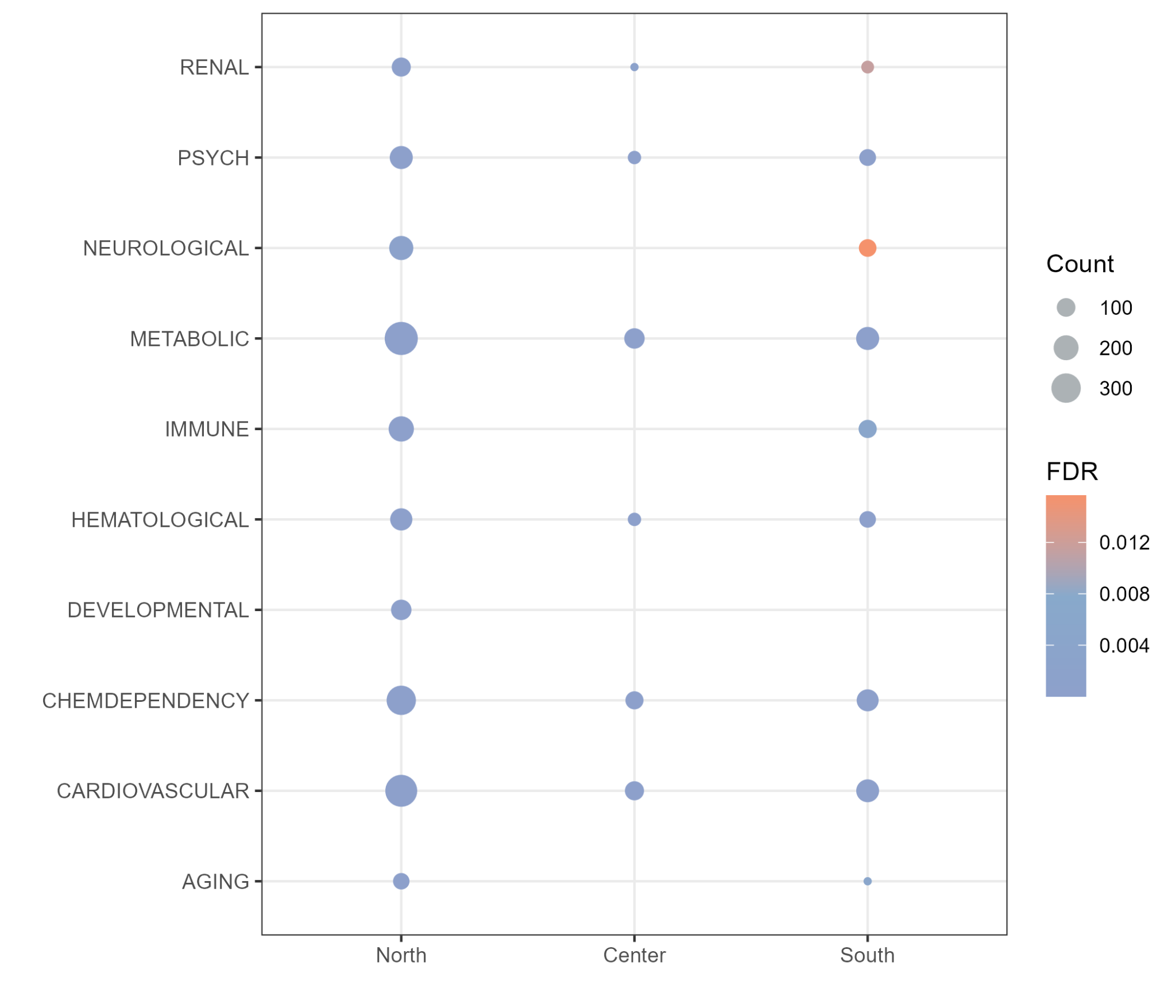

**Figure S10**. Gene disease association for Northern, Center, and Southern Mexican Indigenous Populations with DAVID. Terms represented in the analysis and gene count for each are depicted.

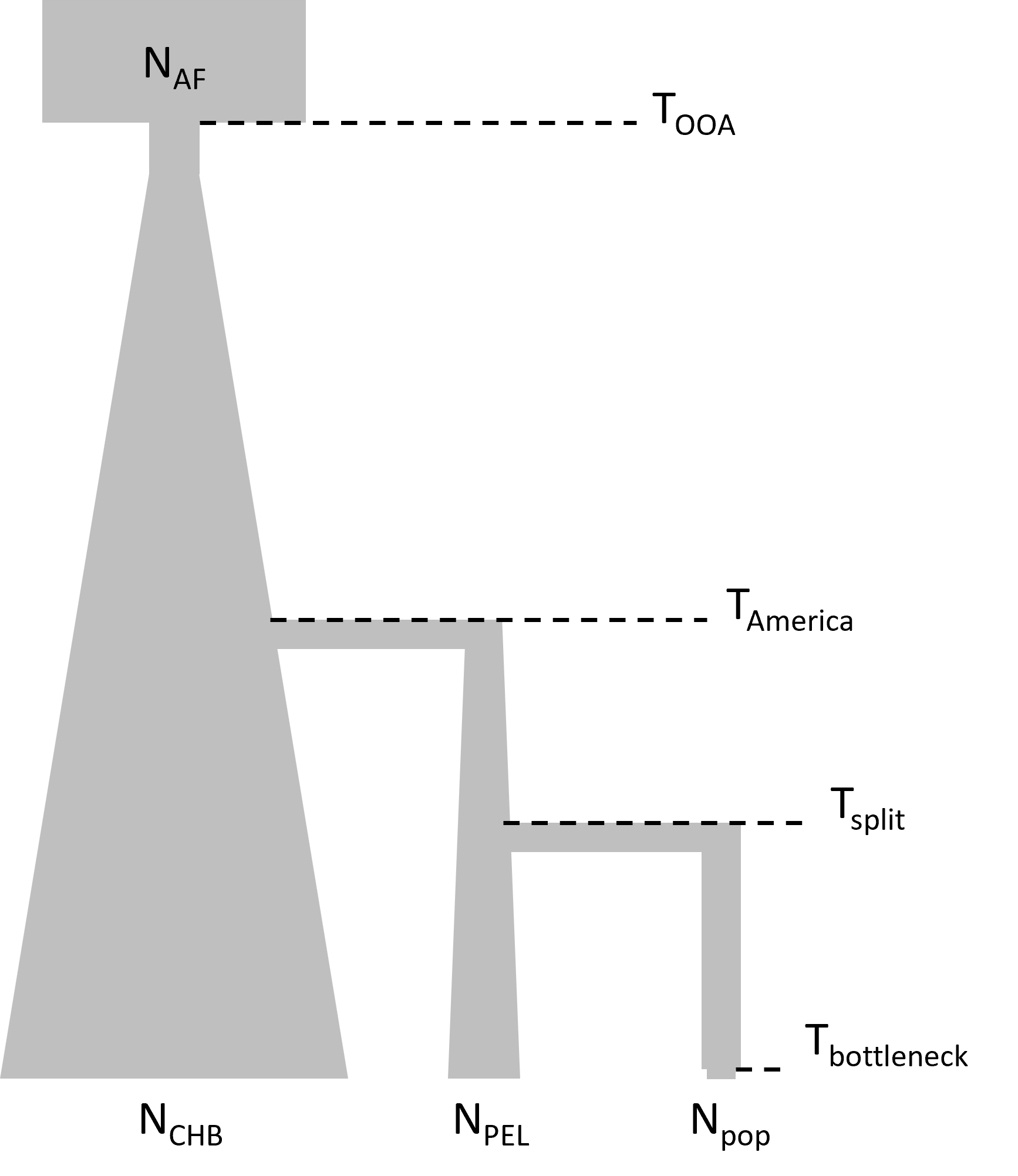

**Figure S11.** Demographic model for Mexican Indigenous Populations. Five demographic events were included. Time is given in generations.
